# Yeast as a platform to dissect *Magnaporthe oryaze* poly(ADP-ribose) polymerase function and evaluate PARP inhibitors

**DOI:** 10.64898/2025.12.29.696956

**Authors:** Rachel E. Kalicharan, Nalleli Payne, Jessie Fernandez

## Abstract

Poly(ADP-ribose) polymerases (PARPs) regulate genome maintenance through NAD^+^-dependent ADP-ribosylation, yet PARP function in fungi remains poorly defined. Here, we reconstituted the activity of the *Magnaporthe oryzae* PARP1 homolog (MoPARP1) in *Saccharomyces cerevisiae*, a genetically tractable organism that lacks endogenous PARP enzymes. Galactose-induced expression of MoPARP1 reduced yeast growth, whereas a catalytically inactive mutant showed no defect, indicating that the growth phenotype depends on PARP catalytic activity. In a multidrug transporter-deficient background, the PARP inhibitor 3-aminobenzamide and the clinically used PARP inhibitor olaparib rescued the growth of MoPARP1-expressing strains, establishing a framework for inhibitor testing *in vivo*. Finally, MoPARP1-GFP localized to the nucleus independent of catalytic activity, supporting correct targeting in this heterologous system. Together, these findings establish yeast as a platform to dissect fungal PARP biology and evaluate chemical inhibition.

## Introduction

Poly(ADP-ribose) polymerases (PARPs) are a large and diverse family of ADP-ribosyltransferases (ARTs) found across eukaryotes. Using nicotinamide adenine dinucleotide (NAD^+^) as a substrate, PARPs transfer ADP-ribose units onto target proteins to generate either mono-ADP-ribosylation (MARylation) or poly(ADP-ribose) (PAR) chains (1-4). In humans, at least 17 PARP family members have been identified. PARP1, the most abundant and best-studied family member, catalyzes the majority of DNA damage-induced PAR synthesis and regulates DNA repair, chromatin dynamics, transcription, and cell death (5). Human PARP1 is a modular enzyme containing regulatory motifs and accessory domains, including zinc finger DNA-binding domains, the tryptophan-glycine-arginine (WGR) domain, and a BRCA1 C-terminal (BRCT) domain, together with a conserved catalytic domain characterized by the histidine-tyrosine-glutamate (HYE) motif (3, 6). PAR signals are reversed primarily by poly(ADP-ribose) glycohydrolase (PARG) and can be pharmacologically blocked using PARP inhibitors (6, 7). Disruption of PARP-mediated ADP-ribosylation is associated with neurodegeneration, metabolic dysfunction, inflammatory diseases, and cancer (4, 8). Notably, the finding that PARP1/2 inhibition can selectively kill BRCA-deficient tumor cells led to the development of multiple FDA-approved PARP inhibitors (9-11). These advances underscore the importance of defining PARP enzymology and the mechanisms by which ADP-ribosylation supports cellular homeostasis.

Despite extensive characterization of human PARPs, far less is known about PARP function across other eukaryotic lineages. Plants encode multiple PARP homologs implicated in DNA damage responses and genome maintenance, and they also contribute to broader stress-response signaling (12-14).

Interest in PARP biology has recently expanded to filamentous fungi, where biological roles have been difficult to dissect, in part because few systems enable robust *in vivo* biochemical interrogation. In the rice blast fungus *Magnaporthe oryzae*, MoPARP1-mediated PARylation of 14-3-3 proteins, which regulate appressorium development, mitogen-activated protein kinase (MAPK) signaling, and virulence, implicates ADP-ribosylation in infection-structure differentiation and host invasion (15). Similarly, in *Fusarium oxysporum* f. sp. *niveum*, FonPARP1 has been shown to be an active nuclear PARP whose activity is enhanced by kinase-dependent phosphorylation, and whose substrate-specific PARylation of the protein disulfide isomerase FonPdi1 regulates protein folding, endoplasmic reticulum homeostasis, and pathogenicity (16, 17). These studies position fungal PARPs as virulence-linked regulators, supporting a reductionist approach to evaluating PARP catalytic outputs and chemical sensitivity in a simplified cellular background.

Heterologous gene expression in yeast provides a powerful approach to define PARP function (18). Prior studies have characterized human and plant PARPs in *Saccharomyces cerevisiae* (19-21). Because yeast lacks endogenous PARP genes and does not synthesize PAR, it offers a genetically “clean” background in which all detected ADP-ribosylation is derived from the introduced enzyme. This genetic simplicity, combined with tractable manipulation and the absence of endogenous PARylation machinery, makes yeast a uniquely useful platform for reconstructing and interrogating PARP activities from diverse organisms.

In this study, we reconstitute and characterize the *M. oryzae* PARP1 homolog (MoPARP1) in the PARP-free background of *S. cerevisiae*. We demonstrate that induced expression of MoPARP1 in yeast leads to a strong, catalysis-dependent growth defect and localizes to the nucleus. A catalytic-site mutant abolishes all enzymatic and phenotypic outputs, confirming dependence on the conserved glutamate residue required for ADP-ribosyltransferase activity. Finally, we show that MoPARP1 is inhibited *in vivo* by 3-aminobenzamide and olaparib, establishing a foundation for using yeast as a screening platform for inhibitors targeting fungal PARPs. Together, our results provide the first biochemical reconstruction of fungal PARP activity in yeast and introduce a versatile system for functional analysis and discovery of antifungal inhibitors.

## Results

### Expression of MoPARP1 reduces yeast growth in a catalytic-dependent manner

To assess whether expression of *Magnaporthe oryzae* PARP1 (MoPARP1) affects yeast growth, we analyzed *Saccharomyces cerevisiae* BY4741 strains carrying an empty vector (EV), wild-type MoPARP1, or a catalytically inactive mutant (MoPARP1-E714A) under repressing (glucose) and inducing (galactose) conditions. Human PARP1 (hsPARP1) was included as a reference for PARP-dependent growth defects previously described in yeast.

Under repressing conditions, all strains displayed comparable growth across serial dilutions, indicating that basal expression did not measurably impact growth (Figure 1A).

**Figure 1.**
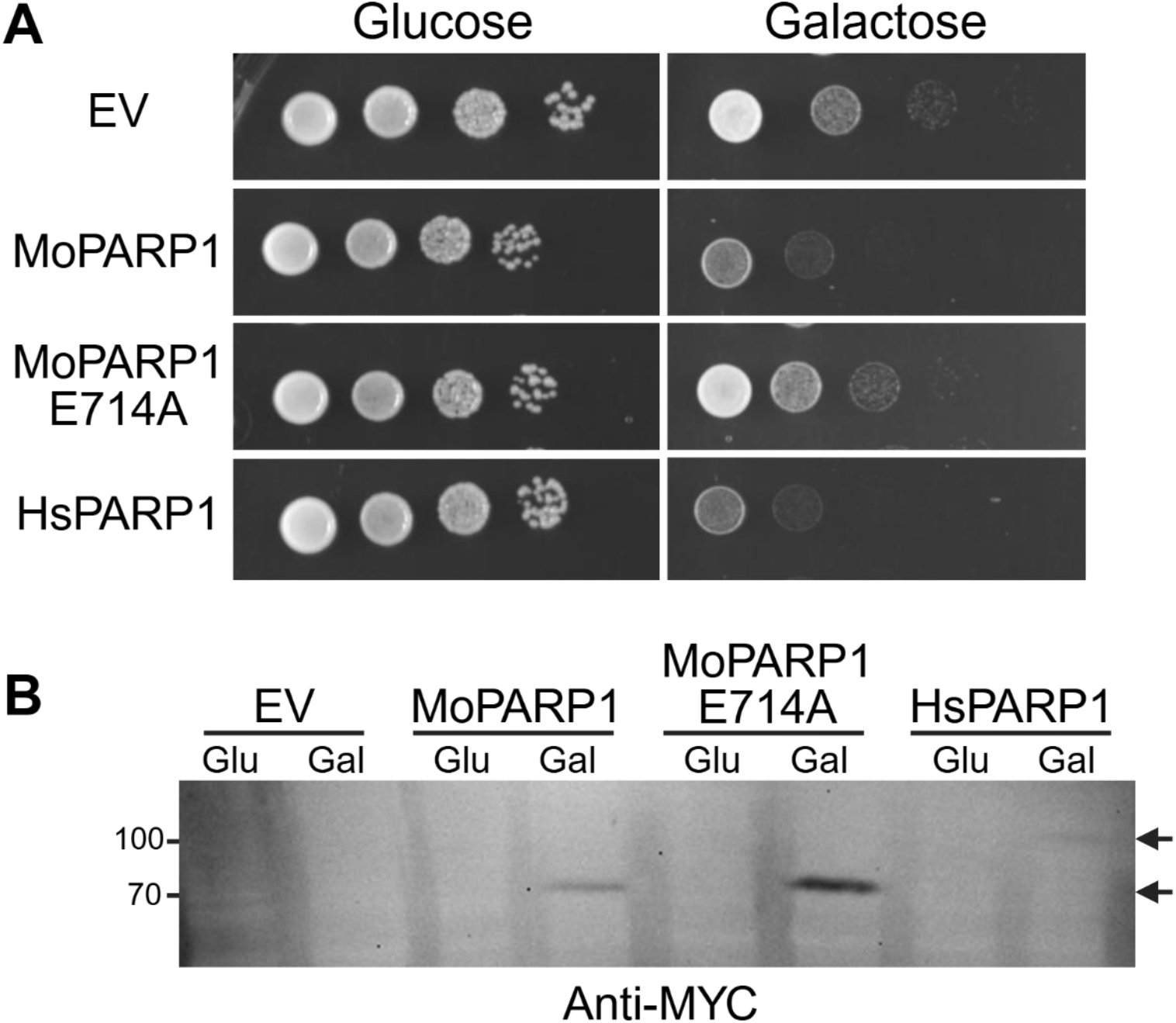
Galactose-inducible expression of PARP1 fusion proteins in yeast and verification of expression. (A) BY4741 yeast strains expressing the empty vector (EV) or indicated PARP1-myc fusion constructs were grown overnight in repressing medium (2% glucose). Cultures were normalized to an OD600 of 1.0, serially diluted 10-fold, and spotted onto plates containing either repressing medium (2% glucose) or inducing medium (2% galactose). Plates were imaged after 2 days of incubation at 30 °C. (B) Induced cultures from (A) were harvested at equal cell equivalents, lysed, and analyzed by SDS-PAGE followed by immunoblotting with an anti-MYC antibody to confirm expression at the expected molecular weights (MoPARP1, 87 kDa; HsPARP1, 110 kDa).

Upon galactose induction, expression of MoPARP1 resulted in a marked reduction in colony formation across the dilutions relative to EV controls. To verify that the observed growth phenotypes were associated with induced PARP expression, we analyzed protein levels by immunoblotting using an anti-MYC antibody. MoPARP1 and MoPARP1-E714A were readily detected at the expected molecular weight in galactose-induced cultures, whereas no signal was observed under repressing conditions or in EV controls (Figure 1B). In contrast, the expression of the catalytic mutant MoPARP1-E714A did not reduce growth, with colony formation comparable to EV on both glucose and galactose media (Figure 1A). These observations indicate that MoPARP1-dependent growth inhibition in yeast requires an intact catalytic residue.

Consistent with prior reports, the induction of hsPARP1 expression resulted in strong growth inhibition on galactose plates (Figure 1A). Notably, MoPARP1 expression produced a qualitatively similar growth phenotype, supporting functional similarity between fungal and human PARP proteins in this heterologous system.

To further characterize growth behavior, we monitored growth dynamics in liquid culture. Under glucose conditions, all strains exhibited similar growth kinetics (Figure 2A, B), consistent with the results of the spotting assay and confirming that the phenotype depends on induced expression. In galactose conditions, strains expressing MoPARP1 or HsPARP1 exhibited a reproducible, pronounced growth delay and achieved lower maximal optical densities compared to EV controls (Figure 2C, D). In contrast, MoPARP1-E714A displayed growth kinetics comparable to EV, consistent with its established effects in yeast. Quantification of growth curves by area under the curve (AUC) analysis supported these observations, with reduced AUC values for MoPARP1- and HsPARP1-expressing strains relative to controls, while MoPARP1-E714A showed no significant difference (Figure 2C, D). Together, these data demonstrate that MoPARP1 impairs yeast growth in a catalytic activity-dependent manner, producing a growth phenotype comparable to that observed upon expression of HsPARP1.

**Figure 2.**
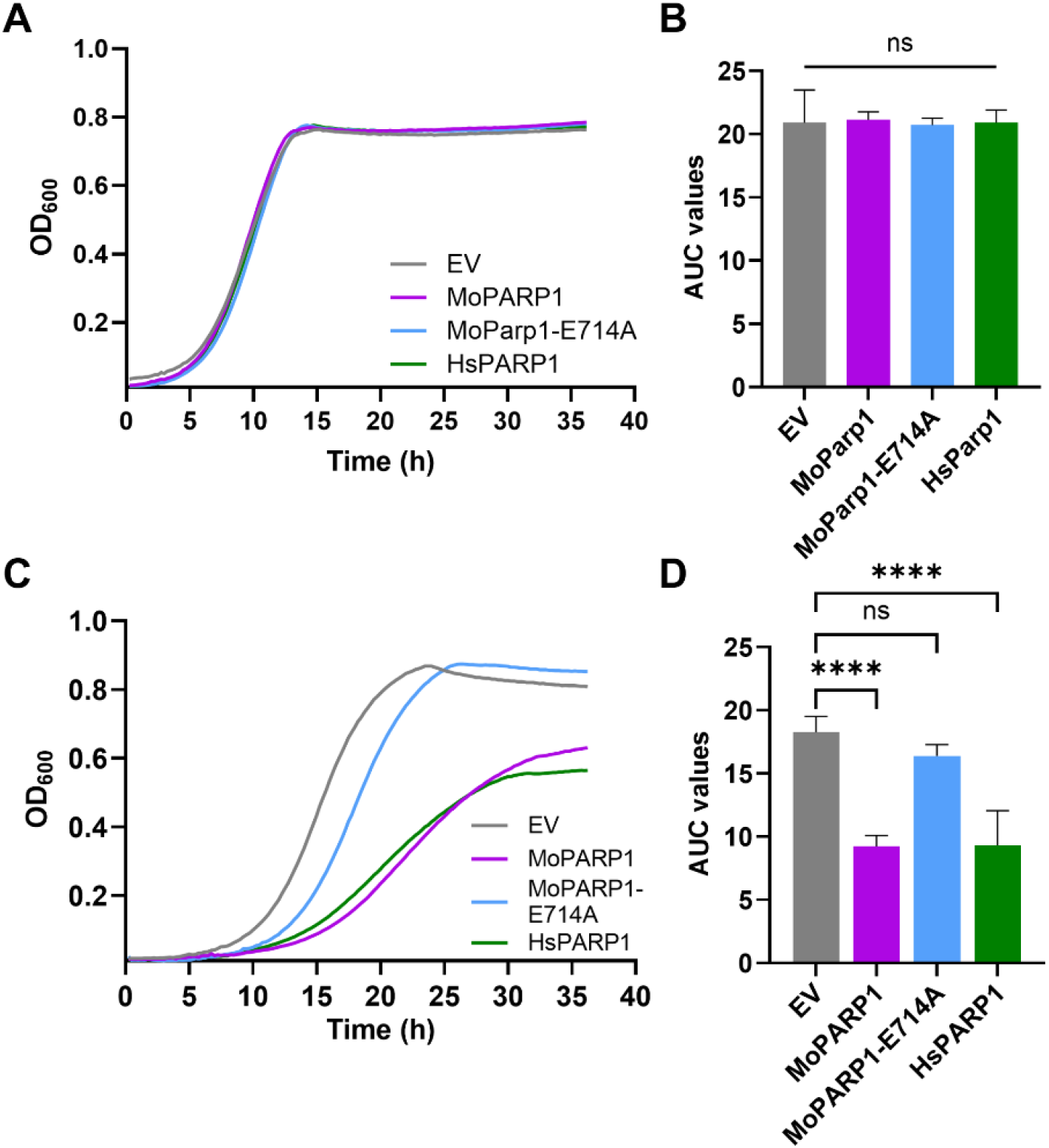
MoPARP1 inhibits yeast growth in a catalytic-dependent manner. (**A**) OD_600_ growth curves of *S. cerevisiae* BY4741 expressing empty vector (EV), MoPARP1, catalytic mutant MoPARP1-E714A, or human PARP1 (HsPARP1) under repressing conditions (glucose). (**B**) Area under the curve (AUC) analysis of (A) shows no significant differences. (**C**) Under inducing conditions (galactose), expression of MoPARP1 and HsPARP1 reduced yeast growth compared with EV, whereas MoPARP1-E714A showed no significant effect. (**D**) AUC quantification of (C). MoPARP1 and HsPARP1 significantly reduced growth relative to EV, while MoPARP1-E714A did not. Data are presented as mean ± SD; statistical significance was determined by one-way ANOVA with Dunnett’s multiple comparisons test; ****, *P* < 0.0001; ns, not significant.

## PARP inhibitor treatment restores MoPARP1-dependent growth inhibition in yeast

To validate whether the growth inhibition associated with MoPARP1 expression is driven by PARP enzymatic activity, we examined the effect of pharmacological PARP inhibition in yeast. These experiments were performed in a BY4741 *pdr5Δ* background, which lacks a major ATP-binding cassette multidrug efflux transporter and thereby enhances intracellular accumulation of small molecules.

As observed in the wild-type background, induction of MoPARP1 expression in *pdr5Δ* cells resulted in strong growth inhibition on galactose-containing media (Figure 3A, B). The addition of the PARP inhibitor 3-aminobenzamide (3-AB; 5 mM), a classical competitive inhibitor of PARP catalytic activity, partially rescued the growth defect of MoPARP1-expressing strains, as evidenced by improved colony formation and increased growth in liquid culture (Figure 3A,B). In contrast, the growth of the strain expressing the catalytically inactive MoPARP1-E714A mutant was unaffected by 3-AB treatment, consistent with the absence of PARP catalytic activity in this variant.

**Figure 3.**
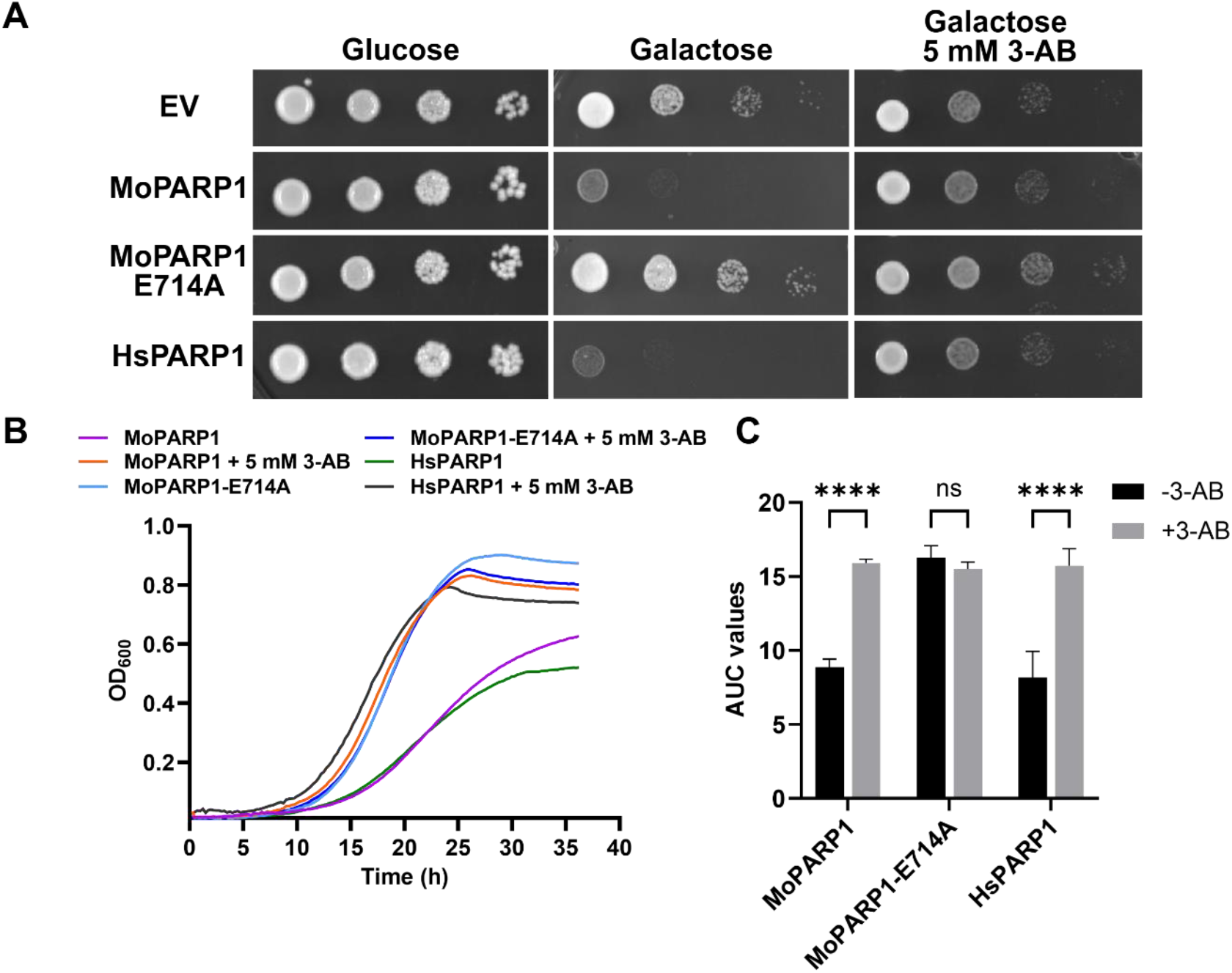
PARP inhibition rescues MoPARP1-mediated growth inhibition under inducing conditions. (**A**) Serial dilution spot assays of BY4741 *pdr5Δ* expressing empty vector (EV), MoPARP1, catalytic mutant MoPARP1-E714A, or HsPARP1. Cells were grown on glucose (repressing) or galactose (inducing) media in the presence of 5 mM 3-aminobenzamide (3-AB). (**B**) Liquid growth curves (OD_600_) under inducing conditions ± 5 mM 3-AB. (**C**) Area under the curve (AUC) quantification of (B). 3-AB significantly increased growth of MoPARP1-expressing cells but had no significant effect on MoPARP1-E714A. Data are presented as mean ± SD. Statistical significance was determined by two-way ANOVA with Šídák’s multiple comparisons test; ****, *P* < 0.0001; ns, not significant.

In line with its established behavior in yeast, the expression of HsPARP1 also caused strong growth inhibition, which was similarly alleviated by 3-AB treatment, serving as a positive control for PARP inhibitor responsiveness (Figure 3A, B). Quantitative analysis of growth curves using AUC measurements confirmed statistically significant restoration of growth for MoPARP1- and HsPARP1-expressing strains upon 3-AB treatment, whereas no significant change was observed for the catalytic mutant (Figure 3B, C).

To extend these findings using a mechanistically distinct inhibitor, we tested olaparib, a clinically used HsPARP1 inhibitor that targets the conserved NAD^+^-binding pocket of the PARP catalytic domain. Under galactose-inducing conditions, treatment with 25 µM olaparib increased the growth of wild-type MoPARP1-expressing yeast in liquid culture relative to untreated controls (Figure 4A, B). This rescue was supported by AUC-based quantification, which showed a significant increase in AUC upon olaparib treatment (*p* < 0.0001). Together with the 3-AB results, these data indicate that MoPARP1-dependent growth inhibition in yeast is enzymatic in nature and can be mitigated by pharmacological PARP inhibition. Collectively, these results demonstrate that MoPARP1-dependent growth inhibition in yeast can be chemically modulated by PARP inhibitors. The concordant inhibitor responses further support functional conservation of key features of the PARP catalytic mechanism and establish yeast as a tractable platform for assessing pharmacological sensitivity of fungal PARP activity.

**Figure 4.**
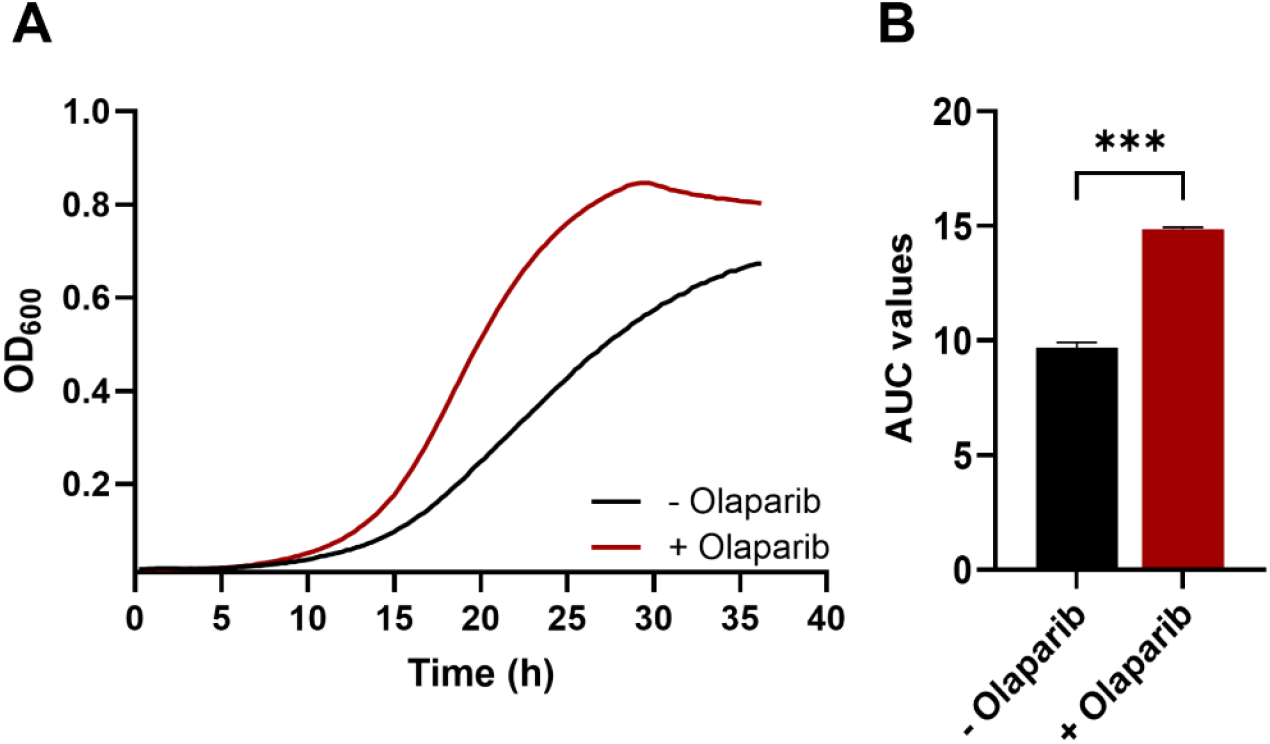
Effect of the PARP inhibitor olaparib on MoPARP1-dependent growth under inducing conditions. Growth of BY4741 *pdr5Δ* cells expressing wild-type MoPARP1 was measured in liquid culture under galactose-inducing conditions in the absence or presence of olaparib (25 µM). Growth was monitored by OD_600_ measurements over time and quantified by area under the curve (AUC) analysis. Data are presented as mean ± SD from three biological replicates. Statistical significance was determined using an unpaired two-tailed t-test with Welch’s correction; ****, *P* < 0.0001; ns, not significant.

### MoPARP1 localizes to the nucleus in yeast independent of its catalytic activity

To validate the yeast heterologous system and determine whether catalytic activity influences subcellular localization, we examined the localization of MoPARP1 in *S. cerevisiae*. Wild-type MoPARP1 and a catalytically inactive mutant, MoPARP1-EA, were expressed as C-terminal GFP fusion proteins from a galactose-inducible vector in BY4741 cells. Fluorescence microscopy showed that both MoPARP1-GFP and MoPARP1-EA-GFP localized exclusively to the nucleus, as indicated by the overlap with Hoechst staining and no detectable cytoplasmic signal (Figure 5). These results confirm correct nuclear targeting of MoPARP1 in yeast and indicate that differences in growth phenotypes between the wild-type and catalytic mutant proteins are not due to altered subcellular localization.

**Figure 5.**
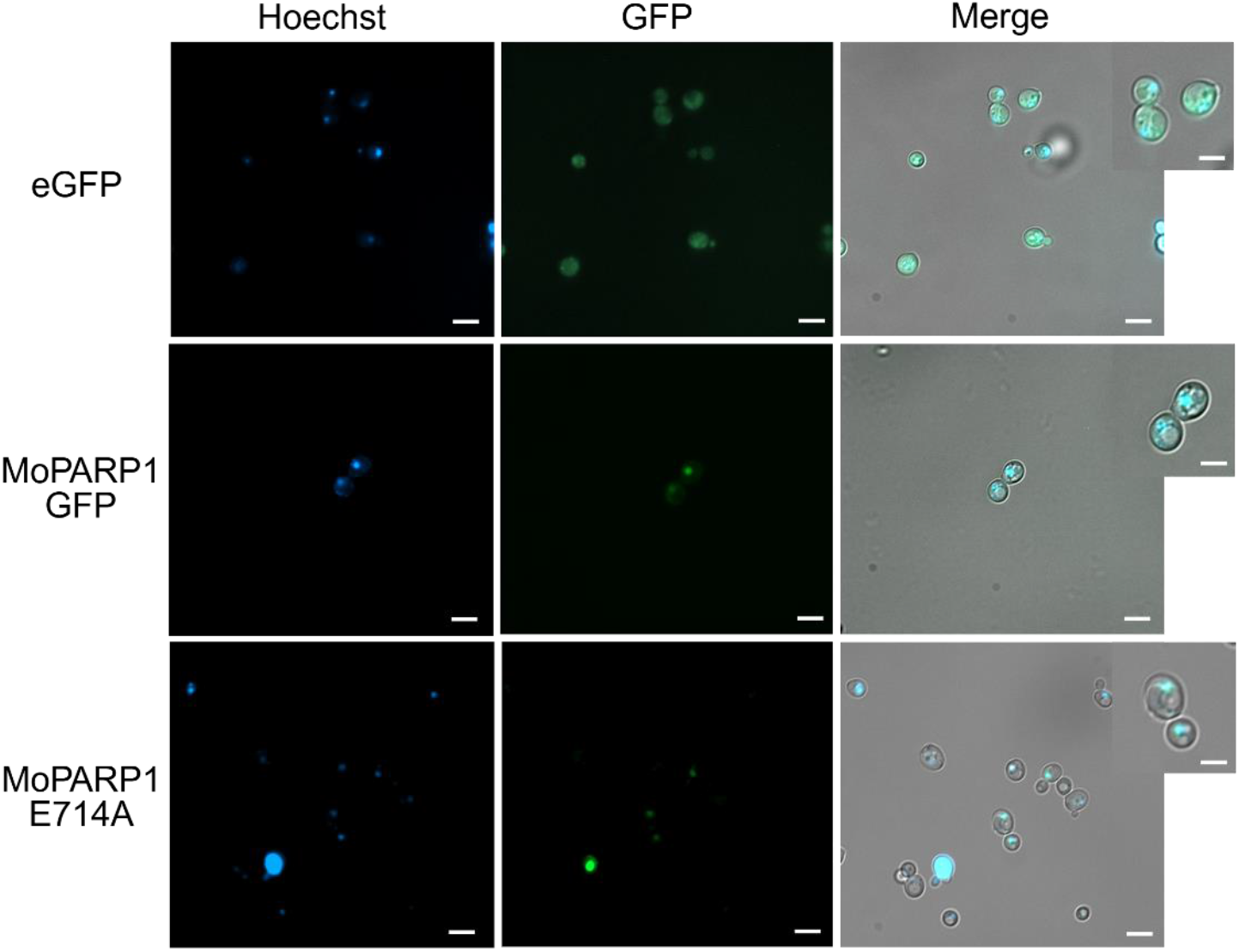
Subcellular localization of galactose-inducible MoPARP1-eGFP fusion proteins in BY4741 yeast cells. BY4741 cells expressing galactose-inducible MoPARP1-eGFP fusion constructs were grown under inducing conditions (galactose) and analyzed by fluorescence microscopy. eGFP fluorescence was used to visualize the subcellular distribution of MoPARP1. Nuclei were stained with Hoechst to serve as a nuclear marker and assess co-localization with MoPARP1-eGFP. Boxed regions indicate areas shown at higher magnification in the inset panels to highlight subcellular localization. Representative fluorescence and corresponding brightfield images are shown. Scale bars = 5 μm

## Discussion

Yeast heterologous expression systems provide a powerful way to dissect protein function in a simplified eukaryotic context, particularly for enzymatic activities that are absent from *S. cerevisiae* and therefore lack confounding endogenous regulation. Because yeast lacks canonical poly(ADP-ribose) polymerases, it provides an essentially background-free platform for isolating PARP-dependent effects and directly connecting catalytic activity to cellular phenotypes (18, 19, 22, 23).

PARylation is a highly dynamic post-translational modification that can rapidly reshape protein interactions, chromatin organization, and stress-responsive signaling. Because PARP activity is tightly coupled to NAD^+^ utilization and generates an amplifying biochemical output, even modest deregulation of PARylation can impose substantial fitness costs. In this context, yeast offers a controlled eukaryotic system in which PARP activity can be examined independently of organism-specific developmental programs or signaling networks. Here, we leverage this system to evaluate the cellular consequences of induced *M. oryzae* PARP1 expression, focusing on growth behavior, subcellular localization, and pharmacological sensitivity.

Using this approach, expression of wild-type MoPARP1 in yeast wielded a pronounced fitness cost, evident in both spot dilution assays and liquid growth curves, whereas growth under repressing conditions remained unaffected. These observations indicate that MoPARP1 is enzymatically active *in vivo* and that its activity alone is sufficient to disrupt cellular growth in a heterologous eukaryotic system. Comparable growth inhibition phenotypes have been reported upon expression of plant and mammalian PARPs in yeast, supporting the idea that deregulated PARP activity is intrinsically deleterious when uncoupled from native regulatory constraints (19, 24, 25). However, it remains unknown which downstream processes are affected by MoPARP1 activity. Given the link between PARP activity and NAD^+^ consumption, future work should investigate whether the growth defect results from NAD^+^ depletion/metabolic stress, and whether it activates yeast stress-response pathways.

The nuclear localization of MoPARP1 in yeast further supports its functional relevance in this system and is consistent with the established chromatin-associated roles of PARP proteins across eukaryotes. MoPARP1 accumulated predominantly in the nucleus under inducing conditions, indicating that nuclear targeting is preserved in the heterologous background. PARP localization has also been examined in *M. oryzae*, where PARP1-GFP fluorescence was readily detected in the nuclei of hyphal cells, while nuclear localization in three-celled conidia was not clearly observed under the conditions tested (15). The conserved nuclear enrichment observed in hyphae and in yeast supports the view that nuclear targeting is an intrinsic property of MoPARP1 and reinforces the relevance of yeast as a system for assessing MoPARP1 localization and function. It remains unclear whether nuclear localization is required for MoPARP1-mediated toxicity and whether specific chromatin-associated interactions underline the observed growth defects.

Chemical inhibition experiments further support the enzymatic basis of the MoPARP1-dependent growth phenotype. Treatment with the 3-AB and the human PARP inhibitor olaparib alleviated MoPARP1-dependent growth inhibition in a multidrug transporter-deficient (*pdr5Δ*) background, consistent with suppression of PARP catalytic activity. 3-AB is a classical competitive PARP inhibitor that binds the conserved nicotinamide-binding pocket within the NAD^+^ site, thereby blocking ADP-ribosyl transfer. The ability to chemically modulate MoPARP1 activity in yeast further highlights the utility of this system for probing PARP enzymatic function. These inhibitor rescue experiments raise questions about which features of the MoPARP1 catalytic domain govern inhibitor sensitivity and whether other clinically relevant PARP inhibitors (PARPis) show similar activity against fungal PARPs.

PARPis have become an important class of targeted therapeutics, particularly in ovarian cancer, and several agents, including olaparib, niraparib, and rucaparib, are approved by the U.S. FDA for maintenance therapy (9). Olaparib was the first PARPi approved as first-line maintenance monotherapy, based on the phase III trial. Mechanistically, olaparib binds to the conserved NAD^+^-binding pocket of PARP enzymes, inhibiting catalytic PARylation and also stabilizing PARP-DNA complexes, thereby promoting DNA damage repair in cells with PARP deficiency (7). Because olaparib is well studied clinically and mechanistically, it provides a useful pharmacological probe for assessing PARP-dependent phenotypes. In the context of this study, the responsiveness of MoPARP1 to olaparib further supports conservation of key catalytic features and validates the use of yeast as a platform for probing fungal PARP activity.

More broadly, yeast has been widely used as a platform to evaluate PARP inhibitor sensitivity, define structure-function relationships, and prioritize small molecules based on phenotypic rescue (19, 23, 26). In this context, MoPARP1-dependent growth inhibition provides a clear and tractable readout that can be leveraged to assess inhibitor responsiveness and guide downstream validation in *M. oryzae*. While compounds identified in yeast would require further characterization for specificity, target engagement, and efficacy in the native pathogen, this approach establishes a foundation for developing chemical probes to interrogate fungal PARP function and explore PARP inhibition as a means to modulate fungal stress responses relevant to pathogenic fitness.

## Material and Methods

### Cloning and vectors

The coding sequence of *Magnaporthe oryzae* PARP1 (MoPARP1) was amplified from *M. oryzae* cDNA using Q5 High-Fidelity DNA Polymerase (New England Biolabs). PCR reactions were set up in 25 µL volumes containing 1× Q5 reaction buffer, 200 µM of each dNTP, 0.5 µM of each forward and reverse primer, and approximately 50 ng of template cDNA. A typical cycling program consisted of an initial denaturation at 98 °C for 30 s, followed by 30 cycles of 98 °C for 10 s, 55-60 °C for 20 s, and 72 °C for 1-2 min, depending on amplicon length, with a final extension at 72 °C for 5 min. The catalytic mutant MoPARP1-E714A was generated by site-directed mutagenesis with Q5 using overlapping primers carrying the desired glutamate-to-alanine substitution at position 714; after amplification, parental template DNA was digested with DpnI at 37 °C for 1-2 h. Human PARP1 coding sequence was obtained from Addgene plasmids (HsPARP1, plasmid #111574) and amplified using Q5 with primer pairs designed to introduce overlaps compatible with Gibson assembly into the pESC-Leu vector (Figure S1). All PCR products (MoPARP1, MoPARP1-E714A, and HsPARP1) were purified with a PCR cleanup kit and inserted into the GAL1-driven pESC-Leu backbone using NEBuilder HiFi DNA Assembly Master Mix (New England Biolabs) following the manufacturer’s instructions. For subcellular localization, the MoPARP1 coding sequence was amplified using Q5 and cloned in frame with GFP into the URA3-selectable pD-eGFP vector via Gibson assembly.

All constructs were confirmed by restriction digestion and Sanger sequencing to verify the correct insert sequence and the presence of the E714A mutation.

### Bacterial transformation and plasmid preparation

Recombinant pESC-Leu and pD-eGFP plasmids were propagated in *Escherichia coli* Top10. Chemically competent Top10 cells were transformed by heat shock at 42 °C for 60 seconds, recovered in SOC medium at 37 °C for 1 hour, and then plated on LB agar containing the appropriate antibiotic. Single colonies were grown overnight at 37 °C in 3-5 mL LB medium with antibiotics, and plasmid DNA was purified using spin miniprep kits. Plasmids were screened by diagnostic restriction digestion and then sequenced to confirm the identity and integrity of MoPARP1, MoPARP1-E714A, and HsPARP1 inserts before yeast transformation.

### Yeast strains, media, and transformation

All yeast experiments were performed in the *Saccharomyces cerevisiae* strain BY4741 (MATa *his3Δ1 leu2Δ0 met15Δ0 ura3Δ0*) and in an isogenic BY4741-*pdr5Δ* mutant (YOR153W; Horizon Discovery), which was used to enhance intracellular accumulation of small-molecule inhibitors in uptake-sensitive assays. For selection of pESC-Leu constructs, strains were maintained on synthetic dropout medium lacking leucine (SD-Leu), whereas selection of pD-GFP constructs for localization was carried out on synthetic complete medium lacking uracil (SD-Ura). For repression of GAL1-driven PARP expression, cells were cultured in SD medium supplemented with 2% (w/v) glucose as the carbon source. For induction of PARP expression, glucose was replaced with 2% (w/v) galactose. Cultures were routinely grown at 30 °C with shaking at 200 rpm. Yeast transformations were performed using the lithium acetate/polyethylene glycol method. Briefly, BY4741 or BY4741-*pdr5Δ* cultures were grown to mid-log phase (OD_600_ approximately 0.4-0.6), harvested by centrifugation, washed with sterile water, and resuspended in 100 mM lithium acetate. Aliquots of competent cells were mixed with plasmid DNA (typically 200–500 ng), boiled salmon sperm carrier DNA, lithium acetate, and PEG 3350, incubated at 30 °C for 30 min, and subjected to a heat shock at 42 °C for 15 min. Transformed cells were recovered briefly in nonselective medium when needed and plated on SD-Leu or SD-Ura agar for selection. Colonies were re-streaked and verified by colony PCR where necessary. For all experiments, BY4741 WT and BY4741-*pdr5Δ* were transformed with the empty vector and with each PARP construct.

### Spotting assays on glucose and galactose

The effect of MoPARP1, MoPARP1-E714A, and HsPARP1 expression on yeast growth was assessed by spotting assays under repressing (glucose) and inducing (galactose) conditions. Single colonies from SD-Leu plates were inoculated into 3-5 mL SD-Leu containing 2% glucose and cultured overnight at 30 °C with shaking. Overnight cultures were harvested by centrifugation, washed twice with sterile water to remove residual glucose, and resuspended in sterile water. Cell density was adjusted to OD_600_ = 0.5, and a series of ten-fold serial dilutions was prepared. Aliquots of 3 µL from each dilution were spotted onto SD-Leu plates containing either 2% glucose or 2% galactose. Plates were incubated at 30 °C for 48-72 h, until colonies were clearly visible, and then photographed. Growth patterns were compared between glucose and galactose for each strain to determine whether induction of PARP expression resulted in growth inhibition. Spotting assays were performed in both BY4741 and BY4741-*pdr5Δ* backgrounds using the same constructs and empty vector controls.

### Cell lysis and immunoblotting

For yeast cell lysis, 1 mL of galactose-induced cultures was collected, and cells were harvested by centrifugation. Pellets were resuspended in 100 µL of ice-cold lysis solution (4% v/v 10 N NaOH, 0.5% v/v β-mercaptoethanol) and incubated on ice for 30 min. Lysates were adjusted to pH 9–10 with HCl, and 2× Laemmli sample buffer was added. Proteins were separated by SDS-PAGE and transferred to polyvinylidene difluoride (PVDF) membranes. MoPARP1 expression was detected by immunoblotting using anti-Myc antibody (1:15,000; clone 9E10, SC-40; Santa Cruz Biotechnology).

### Liquid growth curve analysis

To quantify the effect of PARP expression on yeast growth, liquid growth curves were generated in microplate format. Overnight cultures grown in SD-Leu containing glucose were harvested, washed twice with sterile water, and diluted into SD-Leu containing 2% galactose to an initial OD_600_ of 0.1. For microplate assays, 100 µL of each culture was dispensed into wells of a sterile 96-well plate in triplicate for each strain and condition. Plates were sealed with a breathable membrane and incubated at 30 °C in a microplate reader with orbital shaking. Optical density at 600 nm was measured every 15 minutes for 36 hours. Growth curves were generated by plotting OD_600_ versus time. Quantitative analysis of growth curves and area under the curve (AUC) measurements was performed using GraphPad Prism (GraphPad Software, Boston, MA, USA).

### Inhibitor treatment

Pharmacological inhibition of PARP activity *in vivo* was examined using 3-aminobenzamide (3-AB) and olaparib. A concentrated stock solution of 3-AB was prepared fresh in sterile water, typically at 25 mM. Olaparib was prepared as a concentrated stock solution in DMSO and added to cultures at a final concentration of 25 µM. For inhibitor assays, BY4741-*pdr5Δ* strains carrying pESC-Leu empty vector, MoPARP1-WT, MoPARP1-E714A, or HsPARP1 were grown overnight in SD-Leu with glucose, washed, and resuspended in SD-Leu with galactose to induce PARP expression. For 3-AB treatment, liquid cultures were supplemented with 3-AB to a final concentration of 5 mM, with equivalent volumes of solvent added to control cultures. For plate-based assays, serial dilution spotting was performed on SD-Leu agar plates containing 5 mM 3-AB. For liquid growth assays, cultures were grown in SD-Leu + galactose in the presence or absence of the inhibitor, as described above, and the OD_600_ was monitored over time. For olaparib treatment, assays were performed exclusively in liquid culture under inducing conditions. Growth was monitored by OD_600_ over time to generate growth curves, and the AUC for each replicate was calculated in GraphPad Prism (GraphPad Software, Boston, MA, USA) and used for quantitative comparison between inhibitor-treated and untreated cultures within each construct. The extent to which olaparib alleviated PARP-dependent growth inhibition relative to untreated controls was used as a measure of inhibitor sensitivity in the yeast system.

### Fluorescence microscopy for MoPARP1-GFP localization

Subcellular localization of MoPARP1 was examined in BY4741 cells expressing MoPARP1-GFP from the URA3-selectable pD-GFP vector. Transformants were grown overnight in SD-Ura medium containing 2% glucose, then shifted to SD-Ura medium containing 2% galactose to induce expression for 6 h at 30 °C. Cells were collected by gentle centrifugation, washed once with 1× phosphate-buffered saline (PBS), and fixed with 4% paraformaldehyde. Following fixation, cells were washed twice with 1× PBS and resuspended in a small volume of PBS. For nuclear staining, Hoechst (Sigma) was added to the cell suspension to a final concentration of approximately 1 µg/mL and incubated briefly before mounting on microscope slides. Cells were imaged using a fluorescence microscope (Zeiss Axio Observer 7), and representative images were captured. Images were processed using Fiji (ImageJ distribution) (27). Images were created in https://BioRender.com.

### Statistical analysis

All experiments were performed with at least three independent biological replicates. Growth curve data were analyzed using GraphPad Prism (GraphPad Software, Boston, MA, USA). Differences in growth between strains and conditions were evaluated using two-way ANOVA followed by Dunnett’s and Šídák’s multiple-comparisons test. Area under the curve values were calculated using the trapezoidal method and are reported as mean ± SD. *P* values < 0.05 were considered statistically significant.

## Author Contribution

J.F. conceived the study and designed the experiments. J.F., N.P., and R.E.K, conducted the experiments. J.F., N.P., and R.E.K. wrote and revised the final manuscript.

## Acknowledgement

We thank colleagues whose work could not be cited due to space and scope limitations. We also thank the Fernandez laboratory for constructive feedback.

