## Supplemental Table 1 for "Yeast as a platform to dissect *Magnaporthe oryaze* poly(ADP-ribose) polymerase function and evaluate PARP inhibitors"

**Supplementary Table 1**. Oligonucleotide primers used for Gibson assembly cloning.

| Names | Primers Sequence (5’-3’) | Source |
|---|---|---|
| pESC-Leu MoPARP1 | F: gttgatttccgaagaagacctcgagatgccaccccgacgcagc | This study |
| | R: cttagctagccgcggtaccaagcttcttatcgtcgtcatccttgtaatccatcttgacc | This study |
| pESC-Leu MoPARP1-E714A | F: gttgatttccgaagaagacctcgagatgccaccccgacgcagc | This study |
| | R:_cttagctagccgcggtaccaagcttcttatcgtcgtcatccttgtaatccatcttgacc | This study |
| pDGFP-MoPARP1 | F: ggatcgaattgactctagaggatccatgccaccccgacgcagc | This study |
| | R: catgaattcgagctcggtacccatcttgacccgcaacaggtacc | This study |
| pESC-Leu HsPARP1 | F: gttgatttccgaagaagacctcgagatggcggagtcttctgataag | This study |
| | R: cttagctagccgcggtaccaagcttttaccacagggaggtcttaaaattg | This study |
